# Tracking disease resistance deployment in potato breeding by enrichment sequencing

**DOI:** 10.1101/360644

**Authors:** Miles R Armstrong, Jack Vossen, Tze Yin Lim, Ronald C B Hutten, Jianfei Xu, Shona M Strachan, Brian Harrower, Nicolas Champouret, Eleanor M Gilroy, Ingo Hein

## Abstract

Following the molecular characterisation of functional disease resistance genes in recent years, methods to track and verify the integrity of multiple genes in varieties are needed for crop improvement through resistance stacking. Diagnostic resistance gene enrichment sequencing (dRenSeq) enables the high-confidence identification and complete sequence validation of known functional resistance genes in crops. As demonstrated for tetraploid potato varieties, the methodology is more robust and cost-effective in monitoring resistances than whole-genome sequencing and can be used to appraise (trans)gene integrity efficiently. All currently known NB-LRRs effective against viruses, nematodes and the late blight pathogen *Phytophthora infestans* can be tracked with dRenSeq in potato and hitherto unknown polymorphisms have been identified. The methodology provides a means to improve the speed and efficiency of future disease resistance breeding in crops by directing parental and progeny selection towards effective combinations of resistance genes.

## Introduction

To sustain a potential world population of 9.1 billion by 2050, food production is required to increase by up to 70% compared to 2005-2007 levels (*Fao 2009*). Plant pathogens represent a continuous and serious threat towards this goal and significantly reduce crop yields. The arrival into Europe of the oomycete pathogen *Phytophthora infestans*, for example, led to the Irish potato famine in the mid 1840s. The famine resulted in the death of more than a million people due to starvation and forced the emigration of approximately two million people from Ireland following the successive failures of potato crop production(Donnelly, 2001). Late blight remains the most devastating disease of potato and, alongside other pathogens such as cyst nematodes and viruses, continues to threaten global potato production (Birch *et al*., 2012).

The realisation that resistances against pathogens could be introduced into potato cultivars from wild species (Rudorf, Schaper and Ross, 1949) led to the establishment of international germplasm collections. These collection have been used to introgress genes such as *R1-R11* from *Solanum demissum* into varieties to control late blight (Black *et al*., 1953) and are now being systematically explored to identify additional novel resistances (Vossen, Jo and Vosman, 2014; Van Weymers *et al*., 2016). Owing to the advances in genomics and genetics technologies, numerous functional plant nucleotide-binding, leucine-rich-repeat resistance genes (NLRs) have been subsequently cloned that control diverse pathogens such as potato virus X (Bendahmane, Kanyuka and Baulcombe, 1999), potato cyst nematodes (van der Vossen *et al*., 2000) and the late blight pathogen *Phytophthora infestans* (Hein *et al*., 2009). Similar germplasm resources exist for all major crops throughout the world including wheat, rice, and maize (*FAO* - 2010) and are being explored for beneficial genes including NLRs (Kourelis and van der Hoorn, 2018).

It has been estimated that controlling major diseases through, for example, the informed deployment of functional resistance genes, could contribute over 30% towards crop yield whilst reducing the requirements for chemical applications (Gebhardt and Valkonen, 2001).

Such development in agricultural crop production is dependent on the implementation of new breeding tools to select resistant genotypes and deploy them as varieties. This can be particularly challenging where multiple disease resistances are to be combined in order to take advantage of epistatic interactions (Haesaert *et al*., 2015). To this end we have advanced <u>d</u>iagnostic <u>R</u>esistance gene <u>en</u>richment <u>Seq</u>uencing (dRenSeq), as a novel application to expedite the process of identifying and validating the entire sequence integrity of known functional resistance genes in cultivars and breeding lines. The methodology takes advantage of the focused re-sequencing of NLRs through RenSeq (Jupe *et al*., 2013). RenSeq has previously been used for improving genome annotations and genetic mapping of plant NLRs (Jupe *et al*., 2013; Chen *et al*., 2018), the prioritisation of novel NLRs in wild diploid species (Van Weymers *et al*., 2016; Jiang *et al*., 2018), and of candidate NLRs in combination with long-read sequencing technology (Giolai *et al*., 2016; Witek *et al*., 2016).

Here we demonstrate the applicability of dRenSeq to illuminate the presence of functional NLR genes in tetraploid potatoes, the most important non-cereal food crop (Birch *et al*., 2012). DRenSeq methodology is able to identify and validate all currently known NLRs effective against potato virus X, potato cyst nematode *Globodera pallida* and *P. infestans*. Used as a diagnostic tool in polyploid varieties, dRenSeq provides a robust application that can be utilised to study resistances in any crop where NLRs control diseases.

## Results

### dRenSeq identifies functional NLRs and appraises transgene integrity

The efficacy of dRenSeq in accurately identifying known functional NLRs in crops was initially assessed in 11 transgenic lines derived from the potato variety Desiree that is susceptible towards late blight (Zhu *et al*., 2012, 2014). The plants contained combinations of 14 known NLRs effective against *P. infestans* (*Rpi* genes) (**Figure 1**). Twelve genomic DNA samples from the transgenic and untransformed Desiree plants were indexed prior to NLR enrichment and sequenced on a single lane of Illumina MiSeq (Jupe *et al*., 2013). For dRenSeq, only NLR enriched paired-end reads were mapped, without allowing for any high-quality mismatches, to a reference set of functionally validated NLRs, including their 5’ and 3’ flanking regions (Material and Methods). The representation of individual, full-length NLRs was calculated by extracting the sequence coverage of dRenSeq-mapped reads to the reference coding DNA sequence (CDS) (**Table 1**). We define a resistance gene as “present” if 100% of the CDS is represented by dRenSeq-mapped reads.

**Figure 1.**
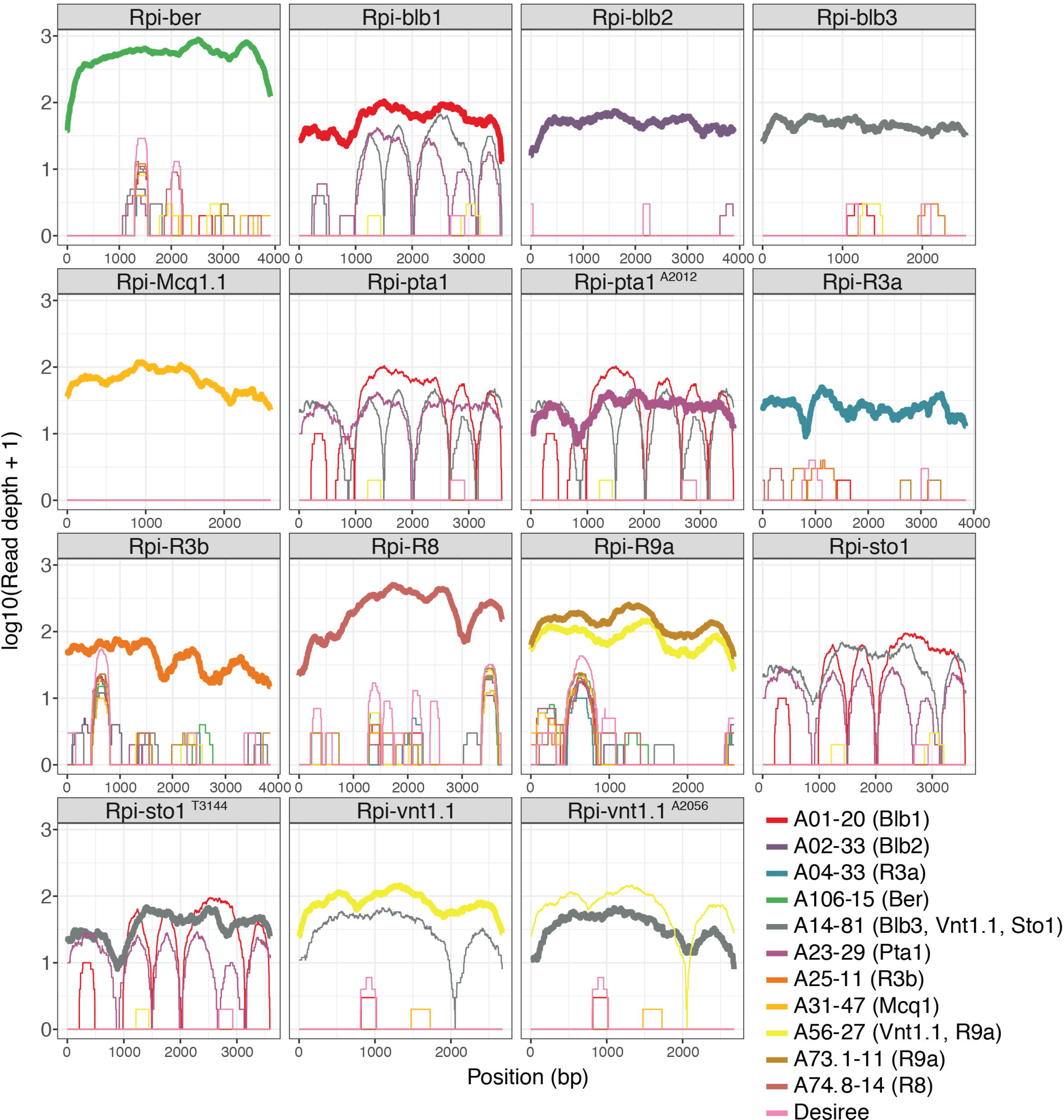
DRenSeq analysis in tetraploid potatoes. DRenSeq analysis of 11 transgenic potato lines derived from the variety Desiree. The sequence representation of known NLRs effective against late blight are shown in each box. The x-axis depicts the coding DNA sequence (CDS) and the y-axis the read-coverage on a log scale. Thick horizontal lines indicate full sequence representation without any sequence polymorphisms between the reference and the NLR enriched reads.

**Table 1.**
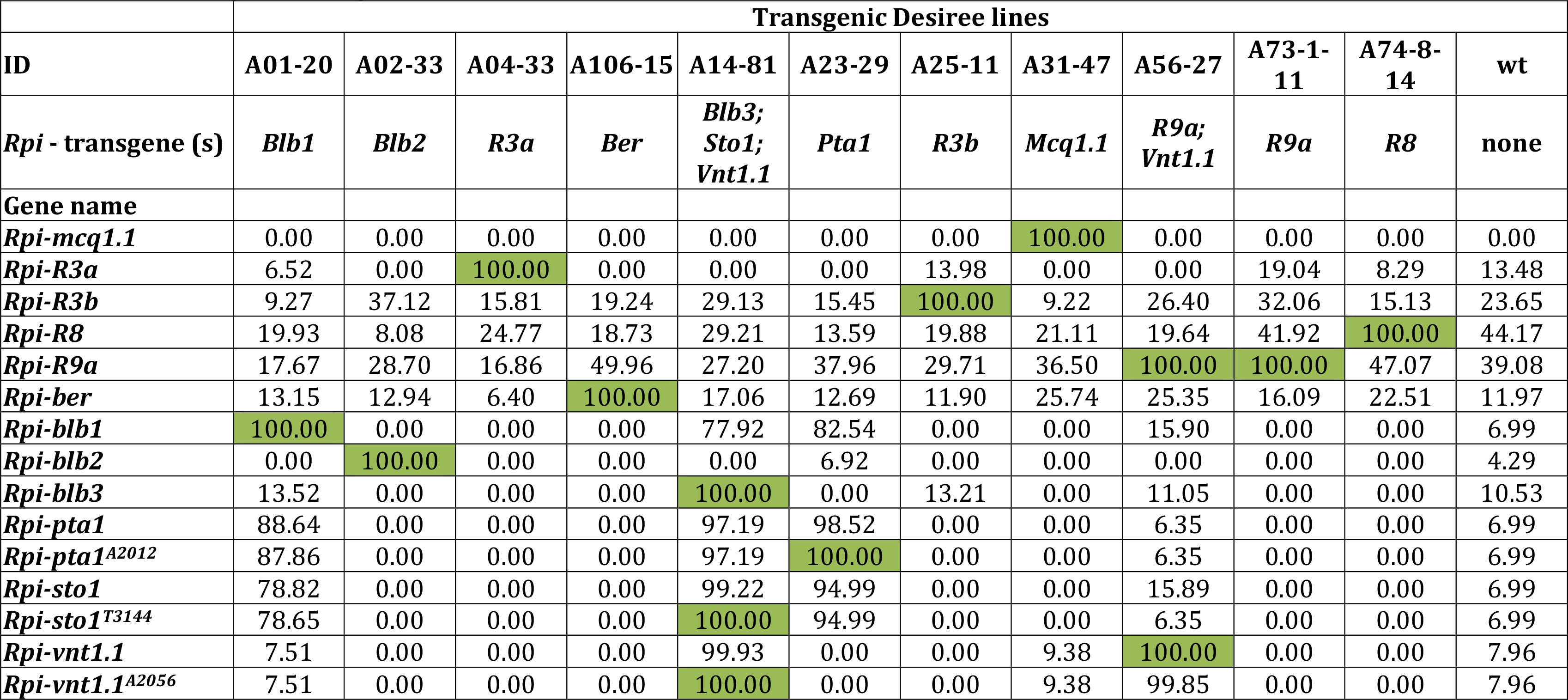
NLR coverage in transgenic Desiree lines. DRenSeq was simultaneously conducted in 11 transgenic Desiree lines alongside a wild-type (wt) Desiree control. The IDs of the transgenic lines and the Resistance to *Phytophthora infestans (Rpi)* nucleotide-binding, leucine-rich-repeat resistances transgenes are shown. The representation of individual, full-length *Rpi* genes was calculated by extracting the sequence coverage of dRenSeq-mapped reads to the reference coding DNA sequence (CDS). Highlighted in green are *Rpi* genes that achieved 100% representation and are therefore classified as ‘present’.

The analysis confirmed the presence of expected single and multiple NLRs but also identified, with high confidence, previously unknown sequence variations within transgenes. Full sequence representation was initially observed for 11 of the 14 NLRs. Surprisingly, *Rpi-sto1* and *Rpi-vnt1.1* in transgenic line A14-81 as well as *Rpi-pta1* in line A23-29 achieved ‘only’ between 98.5% and 99.9% CDS coverage. Remapping NLR-enriched paired-end reads from these lines under less stringent conditions, which allowed a single high-quality SNP per read pair, revealed unexpected sequence variations. We refer to these variants as *Rpi-sto1^T3144^, Rpi-vnt1.1^A2056^* and *Rpi-pta1^A2012^* whereby the affected nucleotide position after the CDS start is indicated alongside the nucleotide substitution (**Supplementary Table 1**). All independent paired-end reads supporting the sequence polymorphisms were monomorphic for the substitutions and no reads were identified that supported the reference nucleotide at these loci (**Supplementary Figure 1a-c**). Crucially, all these predicted sequence polymorphisms were independently validated by Sanger-based re-sequencing of the plasmids used to generate the transgenic lines as well as PCR products derived from the transgenic plants themselves. This shows that the polymorphisms arose during gene cloning and that dRenSeq provides the high resolution required to identify single nucleotide polymorphisms, even amongst a background of related endogenous NLR sequences. Upon generation of new references that incorporate the confirmed polymorphisms, all 14 NLRs were fully represented thus demonstrating the efficacy of the dRenSeq method (**Figure 1 and Table 1**).

### dRenSeq validates *Rpi-vnt1.1* deployment in deregulated GM potato varieties

Following the successful validation of dRenSeq in transgenic Desiree plants, the method was applied to the Innate^®^ Generation 2 transgenic lines Glaciate, Acclimate and Hibernate. These lines have been successfully deregulated in the USA after the introduction of the late blight resistance gene *Rpi-vnt1.1* (Foster *et al*., 2009). Also included in the study were the respective progenitor potato varieties Russet Burbank (for Glaciate), Ranger Russet (for Acclimate) and Atlantic (for Hibernate).

The application of dRenSeq confirmed the presence of functional *Rpi-vnt1.1* in these Innate^®^ transgenic lines and the absence of this gene in all respective progenitor plants (**Figure 2**). In addition, dRenSeq enabled the identification and sequence validation of NLRs effective against diverse pathogens such as nematodes (*Nem*) and viruses (*Virus*) (**Table 2**). For example, dRenSeq revealed that the variety Atlantic contains the late blight resistance gene *R1*, the virus resistance gene *Rx* and the nematode resistance gene *Gpa2*. Within *Gpa2*, a deletion in the intron sequence was identified which does not impact on the NLR protein sequence. We refer to this deletion as to *Nem-Gpa2*^Δ*C2922*^ and subsequently observed the same polymorphism in Hibernate. As anticipated, dRenSeq also independently identified the presence of *Rpi-R1* and *Rx* alongside *Rpi-vnt1.1* in Hibernate, which provides evidence for the robustness of the approach (**Figure 2 and Table 2**). Where we observed significant partial coverage of referenced NLRs other than the transgene *Rpi-vnt1.1* and the closely related gene *Rpi-vnt1.3*, the nucleotide positions of the partial CDS coverage and the depth of the coverage were, overall, in very good agreement between the progenitor lines and the transgenic plants (**Figure 2 and Table 2**). For example, dRenSeq in Russet Burbank revealed partial coverage (42.3%) of *Rpi-R2* that encompassed the 5’ end of the CDS. The equivalent *Rpi-R2* coverage in Glaciate amounted to 42.6% of the CDS with a very similar read-depth and distribution of reads **(Figure 2)**.

**Figure 2.**
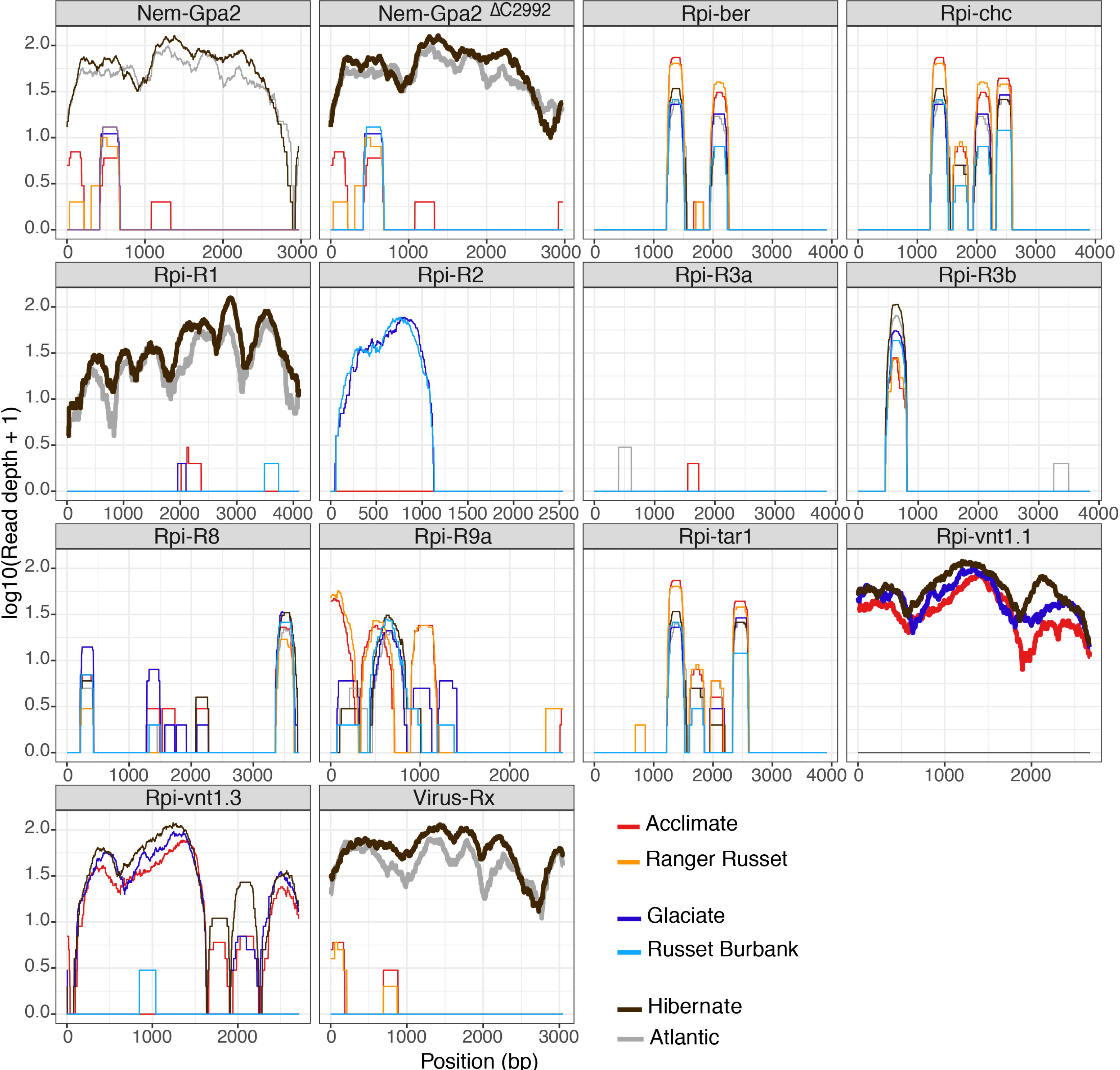
DRenSeq analysis in Innate^®^ generation 2 transgenic lines Glaciate, Acclimate and Hibernate alongside the progenitor varieties Russet Burbank, Ranger Russet and Atlantic. The sequence representation of known NLRs effective against late blight (*Rpi*), nematodes (*Nem*), and viruses (*Virus*) are shown in each box. The x-axis depicts the coding DNA sequence (CDS) and the y-axis the read-coverage on a log scale. Thick horizontal lines indicate full sequence representation without any sequence polymorphisms between the reference and the NLR enriched reads.

**Table 2.** NLR coverage in Innate^®^ generation 2 line. DRenSeq was conducted on Innate^®^ generation 2 transgenic lines Glaciate, Acclimate and Hibernate alongside the progenitor varieties Russet Burbank, Ranger Russet and Atlantic. The name of the varieties and nucleotide-binding, leucine-rich-repeat resistances (NLR) effective against diverse pathogens such as nematodes (*Nem*), late blight (*Rpi*) and viruses (*Virus*) are shown. The representation of individual resistance genes was calculated by extracting the sequence coverage of dRenSeq-mapped reads to the reference coding DNA sequence (CDS). Highlighted in green are resistance genes that achieved 100% representation and are therefore classified as ‘present’.

| ID | Atlantic | Hibernate | Ranger Russet | Acclimate | Russet Burbank | Glaciate |
| --- | --- | --- | --- | --- | --- | --- |
| **Gene name** |  |  |  |  |  |  |
| *Nem-Gpa2* | 99.2 | 98.9 | 18.8 | 24.5 | 8.9 | 8.8 |
| *Nem-Gpa2*<sup>C2922</sup> | 100.0 | 100.0 | 18.8 | 26.4 | 8.9 | 8.8 |
| *Rpi-R1* | 100.0 | 100.0 | 0.0 | 8.6 | 6.1 | 3.5 |
| *Rpi-R1*<sup>T4109</sup> | 100.0 | 100.0 | 0.0 | 8.6 | 6.1 | 3.5 |
| *Rpi-R2* | 0.0 | 0.0 | 0.0 | 0.0 | 42.3 | 42.6 |
| *Rpi-R2-like* | 0.0 | 0.0 | 0.0 | 0.0 | 0.0 | 0.0 |
| *Rpi-R3a* | 5.5 | 0.0 | 0.0 | 4.7 | 0.0 | 0.0 |
| *Rpi-R3b* | 15.7 | 9.4 | 9.1 | 9.2 | 9.4 | 9.3 |
| *Rpi-R3b*<sup>G3111</sup> | 15.7 | 9.4 | 9.1 | 9.2 | 9.4 | 9.3 |
| *Rpi-R3b*<sup>G1696/G3111</sup> | 15.7 | 9.4 | 9.1 | 9.2 | 9.4 | 9.3 |
| *Rpi-R8* | 24.1 | 20.7 | 13.7 | 30.3 | 18.5 | 35.0 |
| *Rpi-R9a* | 27.0 | 30.1 | 46.1 | 39.7 | 39.1 | 42.9 |
| *Rpi-abpt* | 0.0 | 0.0 | 0.0 | 0.0 | 0.0 | 0.0 |
| *Rpi-abpt*<sup>T86</sup> | 0.0 | 0.0 | 0.0 | 0.0 | 0.0 | 0.0 |
| *Rpi-blb1* | 0.0 | 0.0 | 0.0 | 0.0 | 0.0 | 0.0 |
| *Rpi-blb2* | 0.0 | 0.0 | 0.0 | 0.0 | 0.0 | 0.0 |
| *Rpi-blb3* | 0.0 | 0.0 | 0.0 | 0.0 | 0.0 | 0.0 |
| *Rpi-pta1* | 0.0 | 0.0 | 0.0 | 0.0 | 0.0 | 0.0 |
| *Rpi-pta1*<sup>A2012</sup> | 0.0 | 0.0 | 0.0 | 0.0 | 0.0 | 0.0 |
| *Rpi-sto1* | 0.0 | 0.0 | 0.0 | 0.0 | 0.0 | 0.0 |
| *Rpi-sto1*<sup>T3144</sup> | 0.0 | 0.0 | 0.0 | 0.0 | 0.0 | 0.0 |
| *Rpi-vnt1.1* | 0.0 | 100.0 | 0.0 | 100.0 | 7.2 | 100.0 |
| *Rpi-vnt1.1*<sup>A2056</sup> | 0.0 | 96.3 | 0.0 | 96.3 | 7.2 | 98.8 |
| *Rpi-vnt1.3* | 0.0 | 95.5 | 0.0 | 94.2 | 7.1 | 86.9 |
| *Virus-Rx* | 100.0 | 100.0 | 12.8 | 12.4 | 0.0 | 0.0 |

### dRenSeq corroborates NLRs in characterised varieties and identifies sequence polymorphisms

Following the successful validation of dRenSeq in transgenic plants with different progenitors, the method was applied to 12 distinct tetraploid potatoes to further assess the performance of dRenSeq in individual varieties. Similar to the transgenic plants and variety Atlantic, sequence variations compared to reference sequences were identified within certain NLRs and independently detected in multiple varieties (**Supplementary Tables 1, Table 3**). In addition to *Nem-Gpa2*^Δ*C2922*^, polymorphic NLRs *Rpi-abpt^T86^* and *Rpi-R1*^Δ*T4109*^ were identified which contain a synonymous substitution or a deletion in the flanking regions respectively and have no impact on the protein sequence (**Supplementary Table 1**). However, the reference form of *Rpi-R3b* was not observed in any of these varieties whereas *Rpi-R3b* variants with single or double non-synonymous nucleotide substitutions were identified. The single substitution, R3b*G3111*, was present in one variety and the double substitution, *Rpi-R3b^G1696/G3111^*, in 8 varieties including Innovator, Picasso and the late blight differential line 2573 (2). For the differential line 2573 (2), it has been shown previously that *Rpi-R3b* function, such as the recognition of the corresponding effector from *P. infestans, Avr3b*, is maintained (Zhu *et al*., 2014). Similar, *Avr3b* is recognised in Picasso and a strong cell death responds is observed in *Nicotiana benthamiana* upon co-infiltration of *Rpi-R3b^G1696/G3111^* with *Avr3b* (Strachan *et al*., submitted).

**Table 3.** NLR coverage in 12 potato varieties. DRenSeq was conducted on 12 individual potato varieties. The name of the varieties and nucleotide-binding, leucine-rich-repeat resistances (NLR) effective against diverse pathogens such as nematodes (*Nem*), late blight (*Rpi*) and viruses (*Virus*) are shown. The representation of individual resistance genes was calculated by extracting the sequence coverage of dRenSeq-mapped reads to the reference coding DNA sequence (CDS). Highlighted in green are resistance genes that achieved 100% representation and are therefore classified as ‘present’.

| ID | Alouette | Bionica | Cara | Craigs<br>Snow<br>White | Innovator | King<br>Edward | Pentland<br>Ace | Pentland<br>Dell | Picasso | Spunta | Toluca | 2573 (2) |
| --- | --- | --- | --- | --- | --- | --- | --- | --- | --- | --- | --- | --- |
| **Gene name** |  |  |  |  |  |  |  |  |  |  |  |  |
| *Nem-Gpa2* | 29.81 | 21.51 | 98.72 | 27.49 | 18.18 | 58.06 | 8.97 | 30.31 | 97.95 | 0.00 | 0.00 | 7.69 |
| *Nem-Gpa2*<sup>C2922</sup> | 29.81 | 21.51 | 100.00 | 27.49 | 18.18 | 70.70 | 8.97 | 30.31 | 100.00 | 0.00 | 0.00 | 7.69 |
| *Rpi-R1* | 9.34 | 19.21 | 100.00 | 100.00 | 99.20 | 45.73 | 0.00 | 100.00 | 100.00 | 100.00 | 0.00 | 100.00 |
| *Rpi-R1*<sup>T4109</sup> | 9.34 | 19.21 | 100.00 | 99.90 | 100.00 | 45.73 | 0.00 | 99.66 | 99.24 | 99.88 | 0.00 | 100.00 |
| *Rpi-R2* | 0.00 | 58.27 | 0.00 | 9.89 | 50.00 | 7.41 | 8.39 | 58.20 | 4.61 | 0.00 | 0.00 | 57.21 |
| *Rpi-R2-like* | 8.25 | 96.97 | 0.00 | 34.32 | 100.00 | 18.00 | 0.00 | 97.37 | 9.79 | 0.00 | 0.00 | 93.95 |
| *Rpi-R3a* | 100.00 | 100.00 | 100.00 | 0.00 | 100.00 | 63.03 | 100.00 | 100.00 | 100.00 | 6.24 | 17.80 | 100.00 |
| *Rpi-R3b* | 99.12 | 99.66 | 98.94 | 10.93 | 97.72 | 66.07 | 96.99 | 99.92 | 99.33 | 8.54 | 20.12 | 99.12 |
| *Rpi-R3b*<sup>G3111</sup> | 99.43 | 99.79 | 99.07 | 10.93 | 98.65 | 73.16 | 99.22 | 100.00 | 99.66 | 8.54 | 28.79 | 99.64 |
| *Rpi-R3b*<sup>G1696/G3111</sup> | 100.00 | 100.00 | 100.00 | 10.93 | 100.00 | 79.57 | 100.00 | 100.00 | 100.00 | 8.54 | 28.79 | 100.00 |
| *Rpi-R8* | 41.92 | 46.04 | 19.42 | 46.12 | 25.84 | 29.00 | 31.19 | 38.44 | 20.41 | 28.28 | 27.23 | 100.00 |
| *Rpi-R9a* | 32.41 | 50.35 | 27.89 | 47.49 | 5.13 | 50.23 | 47.07 | 40.82 | 36.65 | 39.58 | 51.58 | 100.00 |
| *Rpi-abpt* | 8.27 | 96.14 | 0.00 | 24.86 | 93.77 | 18.05 | 8.39 | 95.82 | 9.81 | 0.00 | 0.00 | 96.30 |
| *Rpi-abpt*<sup>T86</sup> | 8.27 | 100.00 | 0.00 | 24.86 | 97.48 | 18.05 | 8.39 | 100.00 | 9.81 | 0.00 | 0.00 | 100.00 |
| *Rpi-blb1* | 0.00 | 4.12 | 0.00 | 0.00 | 0.00 | 9.10 | 0.00 | 13.89 | 12.14 | 0.00 | 0.00 | 0.00 |
| *Rpi-blb2* | 5.32 | 100.00 | 9.69 | 0.00 | 3.01 | 0.00 | 0.00 | 0.00 | 32.03 | 0.00 | 100.00 | 6.25 |
| *Rpi-blb3* | 0.00 | 55.03 | 0.00 | 9.87 | 44.85 | 0.00 | 0.00 | 50.47 | 9.79 | 0.00 | 0.00 | 44.65 |
| *Rpi-pta1* | 0.00 | 4.12 | 0.00 | 0.00 | 0.00 | 9.11 | 0.00 | 13.90 | 18.63 | 0.00 | 0.00 | 0.00 |
| *Rpi-pta1*<sup>A2012</sup> | 0.00 | 4.12 | 0.00 | 0.00 | 0.00 | 9.11 | 0.00 | 13.90 | 18.63 | 0.00 | 0.00 | 0.00 |
| *Rpi-sto1* | 0.00 | 4.06 | 0.00 | 0.00 | 0.00 | 4.45 | 0.00 | 0.00 | 12.13 | 0.00 | 0.00 | 0.00 |
| *Rpi-sto1*<sup>T3144</sup> | 0.00 | 4.06 | 0.00 | 0.00 | 0.00 | 4.45 | 0.00 | 0.00 | 12.13 | 0.00 | 0.00 | 0.00 |
| *Rpi-vnt1.1* | 95.48 | 21.71 | 0.11 | 8.78 | 9.30 | 0.00 | 7.47 | 8.82 | 7.14 | 6.95 | 0.00 | 0.00 |
| *Rpi-vnt1.1*<sup>A2056</sup> | 88.15 | 21.71 | 0.11 | 8.78 | 9.30 | 0.00 | 7.47 | 8.82 | 7.14 | 6.95 | 0.00 | 0.00 |
| *Rpi-vnt1.3* | 100.00 | 21.38 | 0.11 | 8.65 | 9.16 | 0.00 | 7.36 | 8.68 | 7.03 | 6.84 | 0.00 | 0.00 |
| *Virus-Rx* | 23.51 | 22.20 | 100.00 | 15.89 | 30.77 | 67.16 | 11.69 | 22.82 | 100.00 | 21.87 | 16.16 | 12.35 |

Included in the 12 varieties as additional controls were seven varieties that have been assessed previously for the presence of known NLRs through pathogen assays, gene cloning and/or effector recognition studies (**Supplementary Table 2**). In total, dRenSeq corroborated all 12 previously determined NLRs in the varieties Bionica, Cara, Craigs Snow White, Pentland Ace, Toluca and 2573 (2), which alone contained six known NLRs (**Table 3; Supplementary Table 2**). Interestingly, the variety Cara contains the nematode resistance gene *Gpa2* and the virus resistance *Rx*. This linkage of the 90% identical genes, *Rx* and *Gpa2*, in Cara has been described previously (van der Vossen *et al*., 2000) and the variety was used as a parent in the breeding of Picasso. Both Cara and Picasso share, in addition to *Gpa2*^Δ*C2922*^ and *Rx*, the late blight resistance genes *Rpi-R1, Rpi-R3a, Rpi-R3b^G1696/G3111^*(**Table 3**).

In Pentland Dell, which was released in 1960 as an early example of resistance gene stacking, the presence of late blight resistances *Rpi-R1*, *Rpi-R2* and *Rpi-R3a* had been inferred through the use of differential pathogen isolates^13^. DRenSeq confirmed the presence of *Rpi-R1*, revealed that the *R2*-specific response is based on the presence of the *R2-*family member *Rpi-abpt^T86^*, and that both *R3a* and *R3b^G1696/G3111^* are contained in this variety.

Remarkably, dRenSeq identified the source of resistance in the variety Alouette, which featured on a national list as recent as 2015, as *Rpi-vnt1.3* from *Solanum venturii* (Pel *et al*., 2009). As this is the first example for the deployment of this resistance gene in a commercial variety, the dRenSeq-based inference of *Rpi-vnt1.3* in Alouette was independently confirmed through a) assessing *Avr-vnt1* specific responses in a segregating F1 population and b) PCR-based allele mining **(Supplementary Table 3; Supplementary Figures 2 and 3)**

To generate the segregating F1 population, Alouette was crossed with the susceptible variety Vitalia. In total, 75 progenies from this Al*Vi population were assessed in four replicates for late blight resistances and yielded a 1:1 segregating population with 39 clear resistant and 36 clear susceptible genotypes. This 1:1 segregation ratio provides strong evidence that a single dominant resistant gene is responsible for the late blight resistance in the variety Alouette. From this segregating population 9 resistant and 8 susceptible clones where infiltrated with *Agrobacterium tumefaciens* transiently expressing five individual avirulence genes. Infiltrations were repeated in three plants using three leaves per plant. The scores for hypersensitive responses correlated with the observed disease resistance scores in that all resistant clones yielded an *Avr-vnt1-*specific response, while susceptible plants showed no response. None of the other *Avr* genes tested triggered an HR in any of the tested Al*Vi clones, showing that there was a very specific recognition of *Avr-vnt1* in the resistant plants only (**Supplementary Table 3)**

These data further demonstrate that dRenSeq can be applied to diverse genotypes and is not limited to specific varieties. Critically, where dRenSeq distinguishes different allelic variants, breeders have the opportunity to assess their performance prior to deployment. Functional analysis of allelic variants may distinguish stronger from weaker alleles, but alleles with altered recognition spectra may also be identified.

### dRenSeq-based NLR identification is more cost effective than whole-genome-sequencing

The efficiency of dRenSeq in identifying NLRs was tested in a direct comparison with whole-genome sequencing (WGS) of the potato variety Innovator (**Figure 3**). RenSeq of Innovator yielded 1.7 million high quality MiSeq read pairs representing 778.6Mbp of sequence data which was comparable to all enrichments conducted (**Supplementary Table 4**). DRenSeq analysis revealed the presence of four NLRs; *Rpi-R1*^Δ*T4109*^, *Rpi-R2-like*, *Rpi-R3a* and *Rpi-R3b^G1696/G3111^***(Figure 3; Tables 3 and 4**). It is interesting to note that a distant progenitor of Innovator, AM78-3778, was the source of the molecular characterisation and cloning of *Rpi-R2-like* (Lokossou *et al*., 2009).

**Figure 3.**
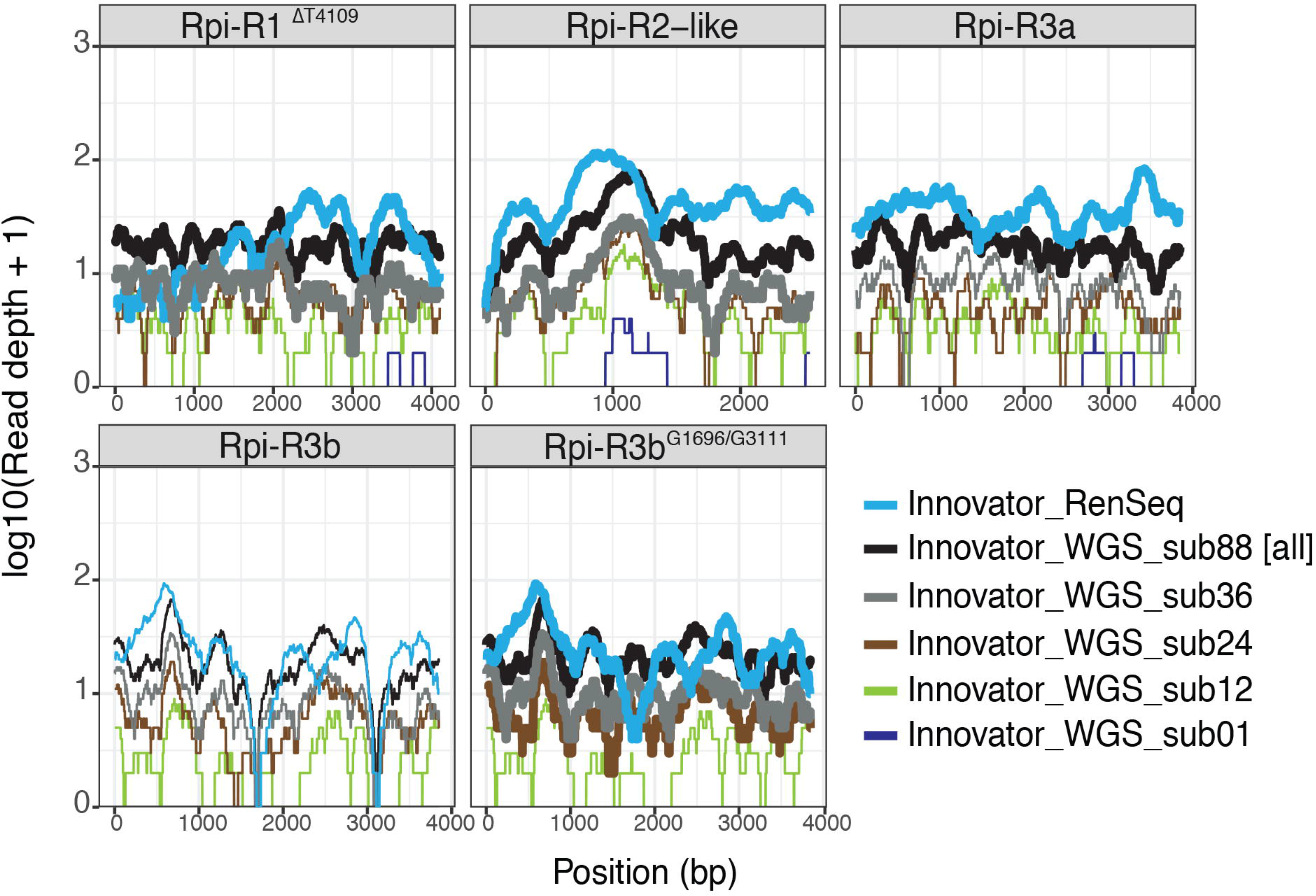
Comparison between dRenSeq and whole-genome shotgun sequencing at different sampling depth for the potato variety Innovator. The sequence representation of NLRs identified in Innovator are shown in each box. The x-axis depicts the coding sequence from start to stop and the y-axis the read-coverage on a log scale. Thick horizontal lines indicate full sequence representation without any sequence polymorphisms between the reference and the NLR enriched reads.

**Table 4.** NLR coverage in commercial potato variety Innovator following RenSeq and whole-genome sequencing. DRenSeq was conducted on potato variety Innovator and compared to whole-genome sequencing (WGS). For the comparison between RenSeq and WGS, subsamples of WGS reads were attained. The sequence volume of WGS reads compared to RenSeq reads are shown in gigabases and x sequence volume [sub 01 = equal amount to RenSeq; sub12 = 12x WGS compared to RenSeq; sub24 = 24x WGS compared to RenSeq; sub36 = 36x WGS compared to RenSeq; sub88 = 88x WGS compared to RenSeq (in this case all WGS data). The IDs of the Resistance to *Phytophthora infestans (Rpi)* nucleotide-binding, leucine-rich-repeat resistances are shown. The representation of individual, full-length *Rpi* genes was calculated by extracting the sequence coverage of dRenSeq-mapped reads to the reference coding DNA sequence (CDS). WGS reads were mapped under the same stringent mapping condition used for dRenSeq. Highlighted in green are *Rpi* genes that achieved 100% representation and are therefore classified as ‘present’.

|  | Innovator |  |  |  |  |  |
| --- | --- | --- | --- | --- | --- | --- |
|  | RenSeq | Whole-genome sequencing |  |  |  |  |
| **Total gigabase pairs** | **0.778** | **69.04** | **28.99** | **19.33** | **9.3** | **0.777** |
| **Gene name** | **RenSeq_all** | **sub88 [all]** | **sub36** | **sub24** | **sub12** | **sub01** |
| *Rpi-R1<sup>ΔT4109</sup>* | 100.00 | 100.00 | 100.00 | 99.34 | 89.76 | 7.36 |
| *Rpi-R2-like* | 100.00 | 100.00 | 100.00 | 95.13 | 93.12 | 20.48 |
| *Rpi-R3a* | 100.00 | 100.00 | 98.29 | 97.04 | 91.22 | 11.33 |
| *Rpi-R3b* | 97.72 | 99.22 | 97.30 | 96.96 | 71.63 | 0.00 |
| *Rpi-R3b<sup>G1696/G3111</sup>* | 100.00 | 100.00 | 100.00 | 100.00 | 83.15 | 0.00 |

WGS of Innovator yielded 228 million high-quality Illumina NextSeq read pairs representing 69,000Mbp of sequence data and thereby over 88x the sequence volume of dRenSeq (**Supplementary Table 4**). After subsampling 778.6Mbp WGS sequences to obtain equal sequence representation compared to RenSeq as well as 12x more WGS sequences, no known NLRs in Innovator were identified with 100% coverage (**Table 4**). Increasing WGS coverage to 24x compared to dRenSeq only identified *Rpi-R3b^G1696/G3111^*, and 36x identified full length *Rpi-R1*^Δ*T4109*^, *Rpi-R2-like and Rpi-R3b^G1696/G3111^* but not *Rpi-R3a*. The NLR *Rpi-R3a* was detected only when all WGS reads were mapped against the references. Importantly, WGS independently confirmed the sequence polymorphisms in *Rpi-R3b^G1696/G3111^* identified by dRenSeq (**Figure 3; Supplementary Figures 4a-b**). Since we routinely combine 12 genomic DNA samples per dRenSeq analysis in a single Illumina MiSeq flow-cell, this demonstrates that RenSeq-based genome reduction is considerably more robust and cost-effective in detecting NLRs than WGS.

## Discussion

Here we demonstrate that dRenSeq enables the parallel identification and sequence validation of multiple functional resistance genes effective against different pathogens. It is currently the only available tool to cost-effectively analyse multiple genotypes in crop breeding programs, identify germplasm with redundant NLRs, and to confirm transgene integrity in commercially available GM crops. The methodology can easily be adapted to include additional functional NLRs, as and when they become available, by ensuring sufficient representation of new genes within the bait library utilised.

A direct comparison between dRenSeq and non-enriched whole-genome sequencing highlights the advantages of dRenSeq. Compared to WGS, only a fraction of the reads is required after NB-LRR gene enrichment to identify and confirm the presence of functional disease resistance genes.

As shown for transgenic Desiree and in other varieties, the sensitivity of dRenSeq, which is achieved through enrichment-based deep-sequencing, is sufficient to determine single sequence polymorphisms in NLRs. This is a prerequisite to certify deployment of functional genes rather than pseudogenised or less effective variants. PCR-based tests are typically unable to identify such variations without sequencing multiple cloned products (Van Weymers *et al*., 2016). Indeed, as recently shown for *R8*, positive PCR amplification does not necessary indicate the presence of functional NLRs as pseudogenised genes can also yield products of similar size (Jiang *et al*., 2018). Therefore, it is often necessary to Sanger sequence PCR products of NLRs, which typically requires the cloning of PCR products and sequencing of recombinant clones. As the average NLR size in potato is estimated to be around 2.7 kb (Jupe *et al*., 2012), at least four Sanger sequencing reactions are required for the analysis of each recombinant clone. Furthermore, as shown for *Rpi-vnt1* in *S. okadae*, numerous recombinants need to be analysed to achieve a true representation of diverse haplotypes (Van Weymers *et al*., 2016). Presuming that the sequencing of five recombinant clones would be sufficient to determine presence/absence and the sequence identity of functional genes, more than 380 Sanger sequencing reactions would be required per single potato variety to assess the 19 genes (**Supplementary Table 5**) that were tested in parallel through dRenSeq (5 clones x 4 sequencing reactions x 19 genes). Since dRenSeq enables the simultaneous analysis and sequence validations of all genes in 12 varieties, the equivalent number of Sanger sequencing reaction would be at least 4,560. Considering the costs and time investments for the PCR analysis and uncertainties in amplifying functional genes from a background of endogenous homologues, dRenSeq represents a cost-effective alternative. Furthermore, a complete dRenSeq analysis from DNA isolation to the computational analysis can be achieved in seven days with little hands on time being required, which again contrasts with the PCR-based alternative detailed above.

Similarly, effector recognition studies are dependent on the discovery of cognate avirulence genes which are not available for all NLRs, are limited by suitable expression systems that work in all host genotypes, and can suffer from a lack of specific plant responses (Vleeshouwers *et al*., 2011). For example, the presence of *Rx* in some varieties including Cara and Picasso precludes the use of the often preferred PVX Agroinfection assay (Du, Rietman and Vleeshouwers, 2014). Similarly, as shown for the cross between Alouette and Vitalia, some plants are more recalcitrant to *A. tumefaciens*-based effector delivery (**Supplementary Table 3**). The observed responses to *Avr-vnt1* and/or *Avr8* co-infiltration with *Rpi-R8* in the cross revealed a spectrum of responsiveness that was measured on a score from 0-2. Therefore, it is necessary to conduct multiple independent inoculations to obtain reproducible results. In addition, these assays typically need special glasshouses, licences and risk assessments.

The ability to track NLRs in crops through dRenSeq is a major advance for modern molecular crop breeding. The methodology can inform on: a) germplasm pedigrees; b) complementary sources for NLR stacking; c) the historic deployment of resistances; d) the geographical differences in NLR deployments. The latter is a prerequisite to gain an insight into the local adaption of pathogens. Indeed, dRenSeq enables the establishment of a reference for already deployed resistance in existing varieties and therefore identify suitable sources of complementary resistance in pre-breeding sources. Thereby, dRenSeq can direct parental selection as well as progeny ranking in crop breeding programs that aim to combine multiple resistances where differential pathogen isolates may not be able to confirm the functionality of the resistances in combination. This could be established for any crop where NLRs control diseases. A prime example is the potential application of a bespoke dRenSeq approach to help control downy mildew of lettuce (*Bremia lactucae*), where a number of functional resistances have been identified already and which could be efficiently combined.

The insight into deployment of NLRs is also relevant for future resistance management strategies. Sensible deployment regulations are required to ensure the durability of resistance in the long run. The dRenSeq application provides a key diagnostic tool to unambiguously determine the NLR contents of new varieties, from which farmers, seed producers, and phytosanitary regulators can make decisions. In the future, similar methodologies could also be adapted for other gene families that control disease including receptor-like kinases and receptor-like proteins once more functional genes have been isolated.

## Material and Methods

### Illumina library preparation

Genomic DNA was extracted from fresh leaf material using the Qiagen DNeasy Plant Mini Kit (Qiagen). The Covaris M220 sonicator (Covaris), was used for the fragmentation of DNA to approximately 500 bp in length, with the following settings: 50W Peak Incident Power, 20% Duty Factor, 200 cycles per burst, 60 seconds treatment time for a 50µL volume with 1µg of starting DNA material. The fragments sizes were checked using a Bioanalyser (Agilent) and no size selection was conducted. Indexed libraries were prepared using the NEBNext library preparation kit for Illumina (NEB). Multiple rounds of AMPure XP bead purification (Beckman Coulter) at a 1:1 ratio of beads to sample was used during the protocol to eliminate fragments smaller than 250 bp in length.

### Targeted enrichment

The genomic DNA libraries were quantified fluorometrically using Qubit (Thermofisher) and 12 indexed libraries were typically pooled prior to enrichment at equimolar amounts so as to achieve 750ng of starting material. The bespoke NB-LRR specific enrichment probe set was purchased from MYcroarray-MYbaits and was based on an improved probe set design described previously (Van Weymers *et al*., 2016). To capture the entire coding DNA sequence (CDS) of known NLRs in the dRenSeq method, the probe set was expanded to include more baits over the individual CDS entireties. The increased bait library [RenSeq library version 4] has been uploaded to http://solanum.hutton.ac.uk. The enrichment protocol was essentially as described for the SureSelect target enrichment system (Agilent) except that human Cot-1 and salmon sperm DNA were omitted from the blocking mix and were replaced by NimbleGen SeqCap EZ Developer Reagent (Roche). Additionally, the blocking mix was supplemented with 1µL of 1000mM universal blocking oligo, containing six inosines in place of the six nucleotide index sequence and a 3’ spacer C3 modification to prevent the oligo from participating in any subsequent PCR amplification. The post capture amplification was performed with the Herculase II polymerase (Agilent). Sequencing was conducted on an Illumina MiSeq platform using the v.2 reagent kit and 2x 250 bp conditions.

### Computational analysis

Illumina reads were trimmed with cutadapt (Martin, 2011) version 1.9.1 to a minimum length of 100bp with a minimum phred quality score of 20 using settings anticipating 3’ anchored adapters. The trimmed reads were mapped to the reference NLRs including 5’ and 3’ flanking regions (**Appendix 1 and 2, Supplementary Table 5**) using bowtie2 (Langmead and Salzberg, 2012) version 2.2.1 in very-sensitive end-to-end mode with discordant read mappings disabled and a maximum insert size of 1000bp. Typically, the score-min parameter was set at L, −0.01, −0.01 which results in a miss-match penalty of five for a 250bp read pair. Consequently, only reads identical to the reference were mapped, except reads containing a variant nucleotide with a quality score <30. For variant discovery, the score-min parameter was set at L, −0.03, −0.03 thus allowing a single high-quality SNP per read pair or a miss-match rate of ~ 0.5%. Due to the synthetic nature of the reference and the high nucleotide similarity of some of the sequences within it, up to 10 mapping positions per read pair were allowed (-k 10). The resulting bam alignments were sorted and indexed using samtools (Li *et al*., 2009) v1.3.1.

Innovator WGS reads were generated by Illumina NextSeq sequencing and were subsampled using seqtk (https://github.com/lh3/seqtk). The read mapping was performed as described above. After mapping, the read coverage and depth of coverage for each reference NLR gene was calculated at each position between the first and last nucleotide of the start and stop codons (Appendix 2) using Bedtools (version 2.25.0) coverage (Quinlan and Hall, 2010). Read depth was log10 transformed and plotted against position using R studio (R Studio Team, 2015) (v1.0.143) and ggplot2. All reads have been submitted to the European Nucleotide Archive (https://www.ebi.ac.uk/ena) with the ENA accession number ERP105478. The reads will be made publicly available on acceptance of the manuscript.

### Segregation of late blight resistance in Alouette*Vitalia population

A cross was made between the late blight resistant potato variety Alouette and the susceptible variety Vitalia. Seeds from this Al*Vi population were sown *in vitro* and 75 germinated seedlings were maintained *in vitro*. Four copies of each genotype were propagated and planted in the late blight trial field in Wageningen. Approximately one month after planting, the field was spray inoculated with a spore suspension of complex *P infestans* isolate IPO-C. From the middle of July disease progress was scored at 4-10 days intervals (**Supplementary Table 3**). An obvious distinction could be made between 39 resistant and 36 susceptible genotypes.

### Agroinfiltration in resistant and susceptible clones of Al*Vi population

Agroinfiltration with *Avr* genes was performed as previously described (Rietman *et al*., 2012). In total 17 randomly selected F1 progeny clones (8 susceptible and 9 resistant) from the Al*Vi population (**Supplementary Table 3**) where infiltrated with *Agrobacterium tumefaciens* transiently expressing five individual avirulence genes. Infiltrations were repeated in three plants using three leaves per plant. The scores for hypersensitive responses were according to Rietman *et al*., (2012) ranging from 0% to 100% cell death, which was converted to a 0-2 scale representing no- to complete cell death in the agroinfiltrated area.

### PCR analysis in resistant and susceptible clones of Al*Vi population

To ascertain which *Rpi-vnt1* gene was responsible for the late blight resistance in Alouette, we conducted PCR reactions with primers LK69 (5’AGCATTGGCCCAATTATCATTAAC3’) and LK70 (5’ATGAATTATTGTGTTTACAAGACTTG3’) on selected clones from the Al*Vi population. The amplification yielded a 1100bp *Rpi-vnt1*-specific amplicon and which was subsequently Sanger sequenced.

---

**Supplementary Figure 1a** Sequence polymorphisms are reliably identified with dRenSeq: Example *Rpi-pta1* in transgenic Desiree line A23-29. Graphical representation of *Rpi-pta1* in transgenic Desiree line A23-29 following dRenSeq-mapping of paired-end reads. The start and stop position of the coding DNA sequence (CDS), relative to the reference sequence that included 5’ and 3’ flanking sequences is shown alongside the position of an intron. Paired-end read mapping was conducted at a 0% mismatch rate, not allowing for any sequence polymorphisms (top), and at a 0.5% mismatch rate, effectively allowing for 1 sequence polymorphism in 200 bp of sequence (middle). The position of a sequence polymorphism relative to the published reference sequence is highlighted. No reads mapped to the polymorphism at a 0% mismatch rate, but numerous paired-end reads with an identical polymorphism (middle and bottom) closed the gap at a 0.5% mismatch rate. The sequence polymorphism was independently confirmed following Sanger-sequencing of plasmids and transgenic lines.

**Supplementary Figure 1b** Sequence polymorphisms are reliably identified with dRenSeq: Example *Rpi-sto1* in transgenic Desiree line A14-81. Graphical representation of *Rpi-sto1* in transgenic Desiree line A14-81 following dRenSeq mapping of paired-end reads. The start and stop position of the coding DNA sequence (CDS), relative to the reference sequence that included 5’ and 3’ flanking sequences is shown alongside the position of an intron. Paired-end read mapping was conducted at a 0% mismatch rate, not allowing for any sequence polymorphisms (top), and at a 0.5% mismatch rate, effectively allowing for 1 sequence polymorphism in 200 bp of sequence (middle). The position of a sequence polymorphism relative to the published reference sequence is highlighted. No reads mapped to the polymorphism at a 0% mismatch rate, but numerous paired-end reads with an identical polymorphism (middle and bottom) closed the gap at a 0.5% mismatch rate. The sequence polymorphism was independently confirmed following Sanger-sequencing of plasmids and transgenic lines.

**Supplementary Figure 1c** Sequence polymorphisms are reliably identified with dRenSeq: Example *Rpi-vnt1.1* in transgenic Desiree line A23-29. Graphical representation of *Rpi-vnt1.1* in transgenic Desiree line A14-81 following dRenSeq mapping of paired-end reads. The start and stop position of the coding DNA sequence (CDS), relative to the reference sequence that included 5’ and 3’ flanking sequences is shown. Paired-end read mapping was conducted at a 0% mismatch rate, not allowing for any sequence polymorphisms (top), and at a 0.5% mismatch rate, effectively allowing for 1 sequence polymorphism in 200 bp of sequence (middle). The position of a sequence polymorphism relative to the published reference sequence is highlighted. No reads mapped to the polymorphism at a 0% mismatch rate, but numerous paired-end reads with an identical polymorphism (middle and bottom) closed the gap at a 0.5% mismatch rate. The sequence polymorphism was independently confirmed following Sanger-sequencing of plasmids and transgenic lines.

**Supplementary Figure 2** A F1 population derived from a cross between varieties Alouette x Vitalia segregates for recognition of *Avr-vnt1*. Agroinfiltration of late blight A*vr* effectors in the Alouette*Vitalia population. **a**: agroinfiltration scheme; **b**: representative pictures of an infiltrated leaf from resistant clone (Al*Vi-59) three days post infiltration. Agroinfiltration with five *Avr* genes was performed in 17 randomly selected clones (8 susceptible and 9 resistant) from the Al*Vi population (Supplementary Table 4). Infiltrations were repeated in three plants using three leaves per plant. In nearly all infiltrated leaves, the positive control (R8+Avr8) showed an obvious hypersensitive response (HR). The 9 resistant clones from the Al*Vi population responded to *Avr-vnt1* infiltration with an HR, while the 8 susceptible genotypes showed no response. None of the other *Avr* genes tested triggered an HR in any of the tested Al*Vi clones, showing that there was a very specific recognition of *Avr-vnt1* only in the resistant plants.

**Supplementary Figure 3** *Rpi-vnt1* PCR analysis in the Al*Vi population. To identify which allele of *Rpi-vnt1* was present in Alouette, we sequenced the PCR products F02 and F03 from clone Al*Vi-4 and F04 and F05 from clone Al*Vi-47 using Sanger sequencing. F02 and F04 were sequenced with primer LK69, and F03 and F05 were sequenced with primer LK70. Sequence alignments show that these amplicons contain a sequence that was identical to *Rpi-vnt1.3*. Only the sequences matching the 5’ end of *Rpi-vnt1* are shown. No polymorphisms were present in the remainder of the amplicon sequences.

**Supplementary Figure 4a**Sequence polymorphisms are reliably identified with dRenSeq: Example *Rpi-R3b* in potato variety Innovator. Graphical representation of *Rpi-R3b* in potato variety Innovator following dRenSeq mapping of paired-end reads. The start and stop position of the coding DNA sequence (CDS), relative to the reference sequence that included 5’ and 3’ flanking sequences is shown. Paired-end read mapping was conducted at a 0% mismatch rate, not allowing for any sequence polymorphisms (top), and at a 0.5% mismatch rate, effectively allowing for 1 sequence polymorphism in 200 bp of sequence (middle). The positions of two sequence polymorphism relative to the published reference sequence is highlighted. No reads mapped to the polymorphism at a 0% mismatch rate, but numerous pairedend reads with an identical polymorphism (middle and bottom) closed the gap at a 0.5% mismatch rate.

**Supplementary Figure 4b** Sequence polymorphisms identified by dRenSeq are also found by whole-genome sequencing (WGS): Example *Rpi-R3b* in potato variety Innovator. Graphical representation of *Rpi-R3b* in potato variety Innovator following mapping of whole-genome sequencing-derived paired-end reads. The start and stop position of the coding DNA sequence (CDS), relative to the reference sequence that included 5’ and 3’ flanking sequences is shown. Paired-end read mapping was conducted at a 0% mismatch rate, not allowing for any sequence polymorphisms (top), and at a 0.5% mismatch rate, effectively allowing for 1 sequence polymorphism in 200 bp of sequence (middle). The positions of two sequence polymorphism relative to the published reference sequence is highlighted. No reads mapped to the polymorphism at a 0% mismatch rate, but numerous paired-end reads with an identical polymorphism (middle and bottom) closed the gap at a 0.5% mismatch rate.

**Supplementary Table 1** Sequence variations identified in resistance genes. Sequence variations in reference genes where identified following dRenSeq mapping of paired-end reads to NLR reference sequences (Appendix 1). Paired-end read mapping was conducted at a 0% mismatch rate, not allowing for any sequence polymorphisms and at a 0.5% mismatch rate, effectively allowing for one sequence polymorphism in 200 bp of sequence. The position of sequence polymorphisms relative to the reference sequence (including flanking sequences) is shown (Pos.). The same polymorphisms are shown in relation to the first nucleotide of the start codon (ATG+) for the coding DNA sequence (CDS) (Appendix 2). The published reference (Ref.) is compared to the variation identified (Var.) Variations include nucleotide substitutions and deletions (-del). If applicable, the nucleotide substitution is shown in the form of a codon change (Codon) and the nature of the change is indicated (NS: non-synonymous; S: synonymous). The resulting amino acid substitution, where applicable, is highlighted.

**Supplementary Table 2** Previously characterised potato varieties/pre-breeding clones confirmed by dRenSeq analysis. Detailed are the names of the potato varieties analysed in this study and used as controls. Previous studies in these varieties have identified NBLRR (Pred. NB-LRR) which were confirmed by dRenSeq (Confirmed NB-LRRs). *Pentland Dell has been described to contain *Rpi-R1, Rpi-R2* and *Rpi-R3* (Malcolmson, 1969). It is now known that *Rpi-abpt* is an *R2* family member (Lokossou *et al*., 2009) and the differentiation of *Rpi-R3* into *Rpi-R3a/R3b* occurred later than the resistance study of Pentland Dell. **This plant was never registered as a cultivar. Hybrid seedlings from crosses between *S. demissum* and *S. tuberosum* were selected from Pentland Field in 1937. An unknown number of backcrosses to *S. tuberosum* resulted in the clone 2573 (2). This plant is used as a late blight differential and pre-breeding clone due to the combination of resistances.

**Supplementary Table 3** A F1 population derived from varieties Alouette x Vitalia segregates for *Rpi-vnt1.3*. Summary of phenotypic and genotypic data for the segregating population derived from crossing resistant potato variety Alouette with susceptible Vitalia. * Disease severity scores in field trial represents the average percentage of blighted foliage (n=4 repeats). S: genotype was found to be susceptible. R: plant was found to be resistant.

** The scores for hypersensitive response were on a scale from 0-2 representing no- to complete cell death in the agroinfiltrated area. r: plant was found to be responsive to Avr-vnt1. n: genotype was found to be non-responsive to Avr-vnt1. Green and yellow highlighted cells represent resistant and susceptible phenotypes, respectively, among plants that have been subjected to the late blight and effector response

**Supplementary Table 4** Illumina sequencing statistics. Shown are the total number of RenSeq enriched and Illumina MiSeq (2×250 bp) generated reads. The sequencing volume in base pairs (bp) is shown for paired-end reads 1 and 2 as well as for the combined sequencing volume. For the comparison between RenSeq and whole-genome sequencing (WGS), subsamples of WGS reads were attained. The relative sequence volume of WGS reads compared to RenSeq reads are shown (x dRenSeq).

**Supplementary Table 5** NLR references. Shown are the gene names, the GenBank ID, and the reference detailing the molecular characterisation of the resistances.

## Appendix 1

FASTA sequence of all reference NLRs used including their 5’ and 3’ flanking region

## Appendix 2

Coordinates of the reference NLR CDS (start – stop)

## Acknowledgement

This work was supported by the Rural & Environment Science & Analytical Services Division of the Scottish Government and the Biotechnology and Biological Sciences Research Council (BBSRC) through awards BB/L008025/1 and BB/K018299/1. We thank Dr Krissana Kowitwanich for the DNA from potato varieties Russet Burbank, Ranger Russet, Atlantic and the transgenic Innate^®^ lines Glaciate, Acclimate and Hibernate.

## Author contributions

I.H., M.R.A., E.M.G, N.C and J.V. conceived the study and wrote the manuscript. M.R.A., S.M.S. and B.H. performed the research. J.V. and R.C.B.H. produced and provided plant material. J.X. performed experiments to show the presence of *Rpi-vnt1.3* in potato variety Alouette. N.C. characterised Innate varieties. T.Y.L and M.R.A. conducted the data analysis. All authors read and approved the manuscript.

## Data availability

All NLR enriched sequence information have been deposited at the European Nucleotide Archive (https://www.ebi.ac.uk/ena) and will be made available upon publication of the manuscript in a peer review journal.

